# Natural killer activation for bladder cancer elimination can be achieved *in vitro* by heat-killed BCG

**DOI:** 10.1101/2020.06.04.129361

**Authors:** Gloria Esteso, Nacho Aguiló, Esther Julián, Omodele Ashiru, Mei. M. Ho, Carlos Martín, Mar Valés-Gómez

**Author notes:** Corresponding author: Mar Valés-Gómez. Department of Immunology and Oncology, National Centre for Biotechnology, CNB-CSIC, Madrid, Spain.

## Abstract

Immunotherapy, via intravesical instillations of *Bacillus Calmette-Guérin* (BCG) is the therapy of choice for patients with high risk non-muscle invasive bladder cancer. The subsequent recruitment of lymphocytes and myeloid cells, as well as the release of cytokines and chemokines, induces a local immune response that contributes to eliminate these tumours. The history of BCG development resulted in a large number of genetically diverse BCG substrains which could stimulate the immune system in different ways. Here, while investigating the capacity of different BCG substrains to promote the activation of NK cells, we confirmed that all the evaluated substrains could activate a cytotoxic CD56^bright^ NK cell population which efficiently degranulated against bladder cancer cells; Tice, Connaught and Moreau were the substrains having a stronger effect. Dead mycobacteria also stimulated PBMC cultures and we demonstrate that subcellular fractions of BCG-Tice could contribute to the induction of this NK cell response. Lipids from BCG-Tice, but not from *Mycobacterium bovis*, stimulated NK cell activation and degranulation, however the aqueous fraction of either bacteria did not activate lymphocytes. Delipidated BCG-Tice activated effector cells (CD3^+^CD56^+^ and NK). These data suggest that different immune subpopulations could be stimulated using different fractions of mycobacteria for cancer elimination.

## INTRODUCTION

Local instillation with the tuberculosis vaccine Bacille Calmette-Guérin (BCG) is the treatment of choice for T1G3 non-muscle invasive bladder cancer (NMIBC) appearing in the form of either papillary tumours or *carcinoma in situ* (CIS). Since the beginning of the use of this therapy several decades ago, the survival time of bladder cancer patients increased notably. However, many questions remain open about the mechanism of action of the BCG against bladder cancer and about the optimal dose and recall instillations to be used in patients.

BCG was generated in 1921, after 13 years of *Mycobacterium bovis* (*M. bovis*) passage *in vitro*, so that this tuberculosis-causing agent acquired a number of mutations and gene modifications, with the loss of the RD1 region being the main cause for attenuation [1]. Through this process, a new strain named BCG was generated. The new cultures of attenuated mycobacteria, perfectly able to reproduce, were distributed worldwide [2,3]. The different host laboratories kept different subcultures of the new bacilli growing. Since no freeze-drying methods were available until the 1960s, BCG acquired new genetic modifications thereby generating independent substrains which received different names, e.g. BCG-Pasteur, -Connaught, -Tokyo, -Danish, -Tice [4-6]. Thus, a whole family of BCG substrains are included under the BCG denomination. Commercial preparations of lyophilized BCG of different substrains are used for intravesical instillations of bladder cancer patients and the viable bacteria content is highly variable ranging from 10^6^-10^9^ [7,4].

BCG has been used for treatment of bladder cancer as intravesical instillations for decades, after the first successful trials in the 1970s [8-10] and FDA approval in 1989 for CIS, followed by an extension in 1998 to treat Ta/T1 papillary tumours. However, because of the way in which BCG was developed and the lack of coordinated clinical trials, many different regimes are used in the treatment of bladder cancer patients with BCG: different hospitals use different substrains, mycobacterial concentrations and schedules [11,12,7]. The general regime of BCG treatment consists of an initial cycle of 6-weekly intravesical instillations, called the induction phase. The main improvement of this therapy during the multiple decades in use, has been the inclusion of a maintenance phase after this initial six-weekly cycle of instillations, consisting in successive cycles of 3 weekly instillations, separated by 3-month rest periods that lasts one to three years. Importantly, clinical studies have directly addressed the effect of the different protocols and, moreover, there are contradicting data on the influence of different substrains of BCG in patient outcome. Clinical trials to study recurrence free survival (RFS) using different substrains are difficult to compare since they include schedules of induction vs induction + maintenance (with different maintenance regimes), one-arm vs two to three-arms, 1 - 5 years follow-up, tens to thousands of patients [12]. In this context, there is a trial including a large number of patients (2,451 patients with primary T1G3 tumors from 23 clinics) in a non-randomized retrospective comparison, finding that BCG Connaught reduced the recurrence rate, overall progression and death compared to BCG-TICE when no maintenance was used, but the opposite was true when maintenance was given. However, for time to progression and overall survival, Connaught and TICE had a similar efficacy [13,14]. These data agree with a prospective randomized single-institution trial with treatment of 142 patients with BCG Connaught or TICE study in which Connaught had better RFS rate. However, the low number of cases did not allow statistical conclusions on progression-free survival, disease-specific survival or overall-survival. In the same report, Rentsch et al [14] treated mice with the same BCG substrains and showed, in the animal model, that BCG Connaught induced a stronger Th1 response, greater priming of BCG-specific CD8^+^ T cells, and more robust T-cell recruitment to the bladder than BCG TICE. Other retrospective studies in BCG-treated bladder cancer patients conclude, however, that there is not much difference between BCG substrains in the treatment of bladder cancer on progression-free survival or disease-specific survival [15].

Since data in the literature point towards a high degree of heterogeneity between the different BCG preparations available for bladder cancer patients, we decided to compare the effect of different BCG substrains on the activation of Peripheral Blood Mononuclear Cells (PBMCs). Both the intensity and the quality of the immune activation might be particular for each BCG preparation. There are data in the literature pointing to a strong Natural Killer (NK) cell component in the mechanism of action of BCG in bladder cancer [16,17]. In fact, we have recently reported, after exposure of PBMCs to BCG, the proliferation and activation of an unconventional cytotoxic subpopulation of CD56^bright^ NK cells that kept functional characteristics of mature NK cells including cytotoxic activity and a high capacity to mediate antibody dependent cellular cytotoxicity (ADCC) [18]. Here, to further characterize the *in vitro* activation of NK cells in response to BCG, we planned a comparison of different substrains in the generation of the anti-tumoral CD56^bright^ NK cell subpopulation. During these studies we discovered that, in addition to different numbers of viable bacilli, the different commercially available presentations of BCG can contain high ratios of dead mycobacteria accompanying the colony-forming units (CFU) and this information cannot be inferred from the supplier leaflet. This finding led us to demonstrate that dead BCG also contribute to the activation of certain pathways of the immune response, in particular, NK cell antitumor degranulation. Evaluating subcellular mycobacterial components from *M. bovis* and BCG-Tice, we determined that fragments from *M. bovis* could strongly provoke lymphocyte proliferation, but less skewed towards and NK cells response when compared with BCG-Tice fragments. Delipidated BCG-Tice was very efficient in stimulating CD56 upregulation, suggesting that non-covalently bound mycobacterial lipids and glycolipids are not strongly involved in NK activation.

## MATERIALS AND METHODS

### Cells, BCG substrains and reagents

Bladder cancer cell lines, T24 and RT112, and the erythroleukemia K562 cell line were previously described [18].

PBMCs from healthy donors (Regional Transfusion Centre, Madrid) were isolated by centrifugation on Ficoll-HyPaque and cultured for one week in RPMI-1640 (Biowest) supplemented with 5% FBS, 5% human male AB serum, 2 mM glutamine, 1 mM sodium pyruvate, 0.1 mM non-essential amino acids, 10 mM Hepes, 100 U/ml penicillin and 100 U/ml streptomycin (Biowest) at a concentration of 1 x 10^6^ cells/ ml in 96-well plates.

World Health Organization Reference Reagents of BCG-Russian (code 07/274) and BCG-Tokyo 172 (code 07/272) substrains were obtained from the National Institute for Biological Standards and Control (NIBSC, UK); BCG-Pasteur strain 1173P2 was originally obtained from Institut Pasteur Paris, BCG-Moreau was a gift from Dr Sylvia Leao, Federal University of Sao Paulo, Brazil); BCG-Tice (OncoTICE) and BCG-Connaught (ImmuCyst) were both commercially available.

### Flow cytometry

For flow cytometry, cells were washed with PBA [PBS supplemented with 0.5% bovine serum albumin (BSA), 1% FBS and 0.1% sodium azide] and incubated with antibodies against surface markers [CD3-CF (Immunostep); CD3-FITC, CD56-PC5, CD16-PE-Cy7, CD4-PC5.5, CD8-PC7 y LAMP1-APC (Biolegend)] at 4°C for 30 min in the dark. For intracellular staining, after surface labelling, cells were fixed with 1% p-formaldehyde for 10 min at RT, permeabilised with 0.2% saponin for 10 min at RT and stained with Perforin-PE, Granzyme B-PE (Biolegend) for 30 min. Cells were washed in PBA and analysed using a Gallios Flow Cytometer (Beckman Coulter). Analysis of the experiments was performed using Kaluza software.

### PBMC stimulation with BCG

Aliquots of reconstituted BCG were prepared in RPMI 10% DMSO and stored at −20°C. For BCG stimulation, experiments were performed as previously described [18]. Briefly, 0.5 x 10^6^ PBMCs/ ml were incubated in 48-well plates with or without BCG at a 1:10 ratio (viable bacteria:PBMC). One week later, cells in suspension were recovered from the co-culture, centrifuged, analysed by flow cytometry.

### Live versus dead mycobacteria counting

Viable bacteria were determined by serial dilutions in Middlebrook 7H10 medium supplemented with Albumin Dextrose Catalase (ADC). To obtain heat-killed BCG, bacteria were washed in PBS and treated at 100°C for 20-30 min. The total number of BCG, including live and dead bacilli, was calculated by flow cytometry using a protocol optimized for this study as follows. A calibration curve was prepared, using a log-phase growth culture of BCG, in Middlebrook 7H9 medium supplemented with ADC, assuming that the whole culture was viable. Then, serial dilutions (1:2) were prepared and analysed by flow cytometry. Bacterial viable units in the culture were calculated by plating in solid medium and counting colonies for each standard curve point. For these analyses, FSC and SSC parameters were set in the flow cytometer to visualize bacteria (Supplementary Figure 1). Samples were acquired, using a Beckman Coulter Gallios flow cytometer, at constant speed during 30 s and the number of events was defined as the total amount of bacteria. For analysis, weasel software was used. Data of the different dilutions were used to build the calibration curve and, the latter to interpolate and calculate the total amount of BCG present in different formulations.

### Degranulation experiments

Untreated or BCG-treated PBMCs were used as effector cells; bladder cancer cell lines or the erythroleukemia K562 (as positive control for NK degranulation) were used as target cells, always pretreated with HP1F7 antibody to block MHC-I. The percentage of NK cells was determined for each donor, so that equivalent effector NK cell ratios were set up for the different donors and conditions. NK cells were allowed to degranulate in co-cultures of PBMCs and targets for 2 h at an E:T ratio of 5:1 [which corresponds usually to 1:2 NK:target ratio]. Surface expression of LAMP1 (CD107a) was analyzed by flow cytometry. Statistical analyses were performed using the Prism 6 software.

### Mycobacterial subcellular fractions

The following reagents were obtained through the NIH Biodefense and Emerging Infections Research Resources Repository, NIAID, NIH: γ-irradiated whole cells *M. bovis*, Strain AF 2122/97 (ATCC^®^ BAA-935^™^) Catalog No. NR-31210; Total Lipids, Catalog No. NR-44100, Cell Membrane Fraction, Catalog No. NR-31214 and Cytosol Fraction, Catalog No. NR-31215 from *M. bovis;* Trehalose-6,6-dimycolate (TDM), Catalog No. NR-14844 and Mycolic Acid, Catalog No. NR-14854 from *M. tuberculosis*, strain H37Rv.

To obtain BCG-Tice fragments, BCG-Tice was grown at 37°C on Middlebrook 7H10 medium supplemented with Oleic-Albumin-Dextrose-Catalase (OADC). After 4 weeks, cells were scraped from the surface and non-covalently attached cell wall lipids were extracted, first with chloroform/methanol (2:1, v/v), and then with chloroform/methanol (1:2, v/v) mixtures. Pooled organic extracts, containing the lipids, were combined, dried and partitioned with chloroform/methanol/water (8:4:2 v/v). Aqueous phase (“aqueous” antigen) and chloroform phase (“lipid” antigen) were then separated and evaporated to dryness. Delipidated bacteria were collected and stored in aliquots in sterile glass tubes (“delipidated” antigen).

## RESULTS

### Differential activation of CD56^bright^ NK cells by BCG substrains depends on the number of total mycobacteria

To test whether the different BCG substrains used in the clinics could differently activate the immune response, experiments to study the variability of mycobacteria in the context of NK activation were initiated. The generation of an anti-tumoral NK cell population by stimulation with BCG characterized by upregulation of the surface marker CD56 on NK cells [18], was reported recently, using the clinical grade preparation OncoTICE^®^ at a ratio of 1:10 (10^5^ live bacteria to 10^6^ PBMC). So firstly, we compared the effect of two reference BCG vaccines with OncoTICE^®^ in the upregulation of the surface marker CD56 on NK cells using the same ratio as previously. Incubation of PBMCs with both Tokyo and Russian substrains (lyophilized preparations obtained from NIBSC) also resulted in an increase in the percentage of CD56^bright^ NK cells (Figure 1A). Because different commercial formulations contain different concentrations of BCG, laboratory grown cultures of the above mentioned different substrains were subsequently used to plan new experiments using equal numbers of mycobacteria. When using laboratory grown substrains, upregulation of CD56 also could be observed, although surprisingly, the percentages of CD56^bright^ cells obtained were much lower with the laboratory preparations in comparison with OncoTICE^®^ (Figure 1B). Since the pellet obtained after centrifugation of the suspensions from commercially available preparations was substantial, we hypothesized that a larger number of bacteria, presumably dead, were present in addition to the live BCG indicated in the leaflet. Thus, the numbers of live *vs* dead BCG were analyzed in each lyophilized sample. Live BCG were measured by counting CFU in Middlebrook 7H10 agar plates and the total number of BCG was estimated by flow cytometry, using the protocol optimized for the present study (see material and methods and Supplementary Figure 1). These numbers were compared with the concentration range provided by the manufacturer. While the CFU range provided in the product insert agreed with the numbers that were obtained by the culture method, the number of total BCG obtained by flow cytometry was, in general, much higher. Table 1 summarizes the numbers obtained for the different BCG preparations. For example, OncoTICE^®^ contains 2-8 x 10^8^ viable BCG per vial, according to the manufacturer’s information but the total number of mycobacteria measured by flow cytometry was 5.95x 10^9^, thus, only around 10% of mycobacteria present in the vial were alive after reconstitution in buffer. Similarly, only 38% of mycobacteria were alive in the BCG Tokyo lyophilised ampoule (NIBSC, WHO reference reagent, Japan BCG laboratory) (Table 1).

**Table 1.** Live *vs* dead BCG in commercial lyophilised preparations.

| SUBSTRAIN | BCG CFU* | TOTAL BCG<sup>#</sup> |
| --- | --- | --- |
| TICE | $2-8 \times 10^8$ (3mg) | $6.0 \times 10^9$ |
| CONNAUGHT | $1.8-15.9 \times 10^8$ (81mg) | $5.2 \times 10^9$ |
| RUSSIAN | $3.4 \times 10^6$ (0.5mg) | $1.5 \times 10^7$ |
| TOKYO | $5 \times 10^7$ (1mg) | $1.3 \times 10^8$ |

\* CFU provided by the manufacturer or NIBSC [19] (expressed as dry weight) and confirmed in the laboratory
<sup>#</sup> Number of BCG cells counted using flow cytometry analysis (see methods).

**Figure 1.**
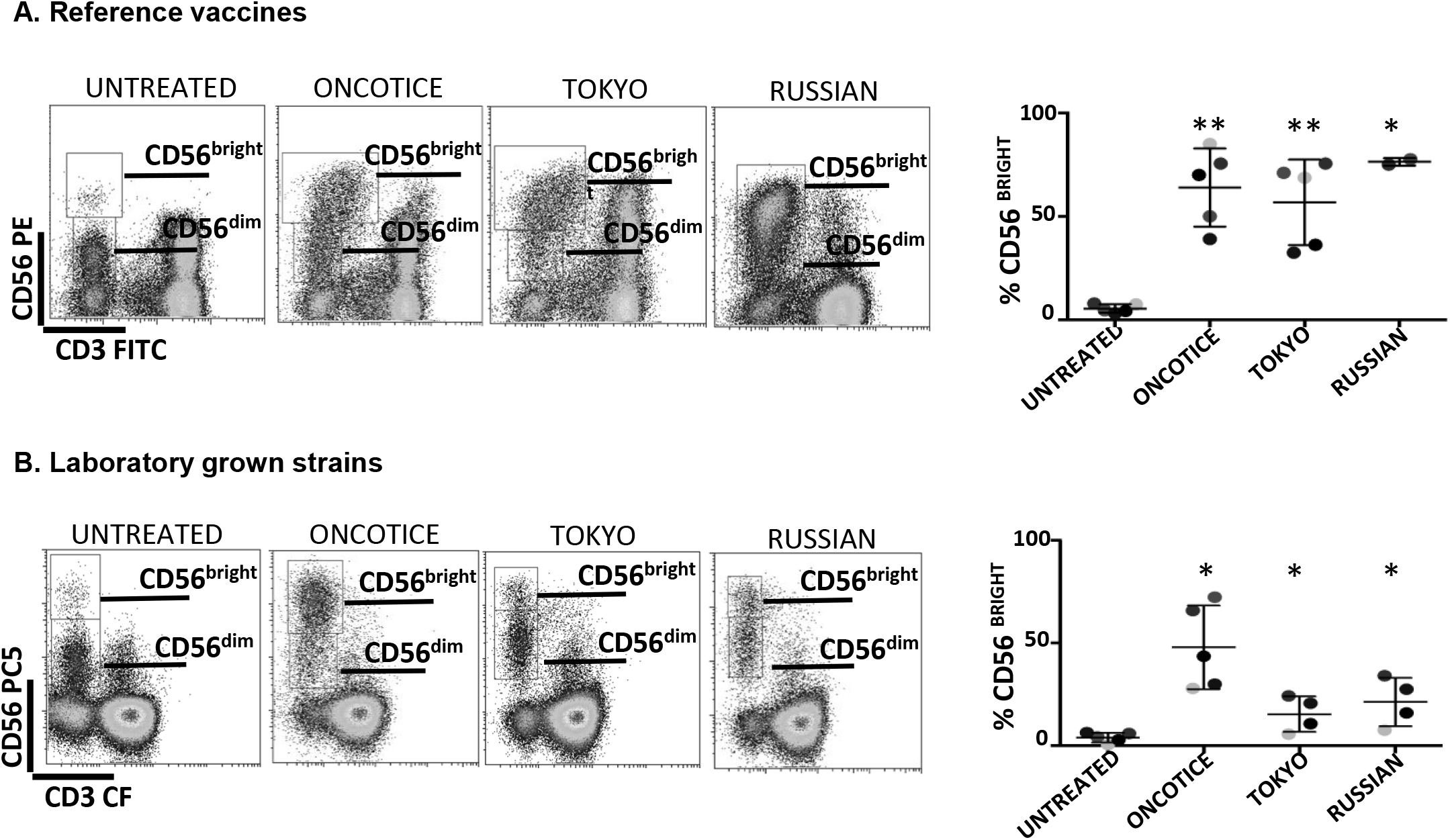
NK cell activation, based on the increase of CD56^bright^ cells, occurs after incubation with several substrains of BCG. PBMCs from healthy donors were incubated with or without the indicated substrains of BCG at a 1:10 ratio (viable bacteria to PBMC). Commercial OncoTice was used for comparison. At day 7, cells in suspension were recovered from the co-culture, centrifuged and analysed by flow cytometry. The percentage of the CD56^bright^ and CD56^dim^ populations within NK cells was obtained for each donor and plotted in a bar graph. Statistical analysis was performed using T-student test to compare each substrain with OncoTice (*, p<0.05; **, p<0.01). **A. Comparison of reference vaccines with Oncotice. B. Comparison of laboratory-grown substrains with Oncotice.**

Thus, the differences observed in NK activation between the commercial lyophilized product (OncoTice) and the laboratory culture preparations might be due to the presence of large numbers of dead bacteria in the former.

### Dead mycobacteria can activate anti-tumour CD56^bright^ NK cells to degranulate against bladder cancer cell lines

Since we hypothesized that dead BCG could contribute to the activation of CD56^bright^ NK cells, the capacities of clinical grade Tice and several laboratory-cultured BCG preparations to stimulate PBMCs were compared using either viable or heat-inactivated BCG. In order to avoid heterogeneity due to culture/ production conditions, the different substrains were grown in laboratory using the same culture medium and method. BCG added to PBMCs were either mostly viable (at log phase), completely inactivated or mixed 1:2. In all the cases, PBMC activation was observed as shown by the emergence of lymphoblast cells in the FCS^high^/SSC^high^ region (Figure 2A, Supplementary Figure 2). The percentage of activated lymphocytes was plotted for the different donors showing very similar percentages numbers with viable and heat-killed bacteria. The commercial product, OncoTICE, was used for comparison, since it was the one used for our initial description of NK cell activation. CD56^bright^ NK cells were present in all the cases (Figure 2B). The percentage of CD56^bright^ NK cells increased significantly with respect to untreated cells even when PBMCs were incubated with 100% heat-inactivated BCG, suggesting that the activation of NK cells does not require interaction with live mycobacteria.

**Figure 2.**
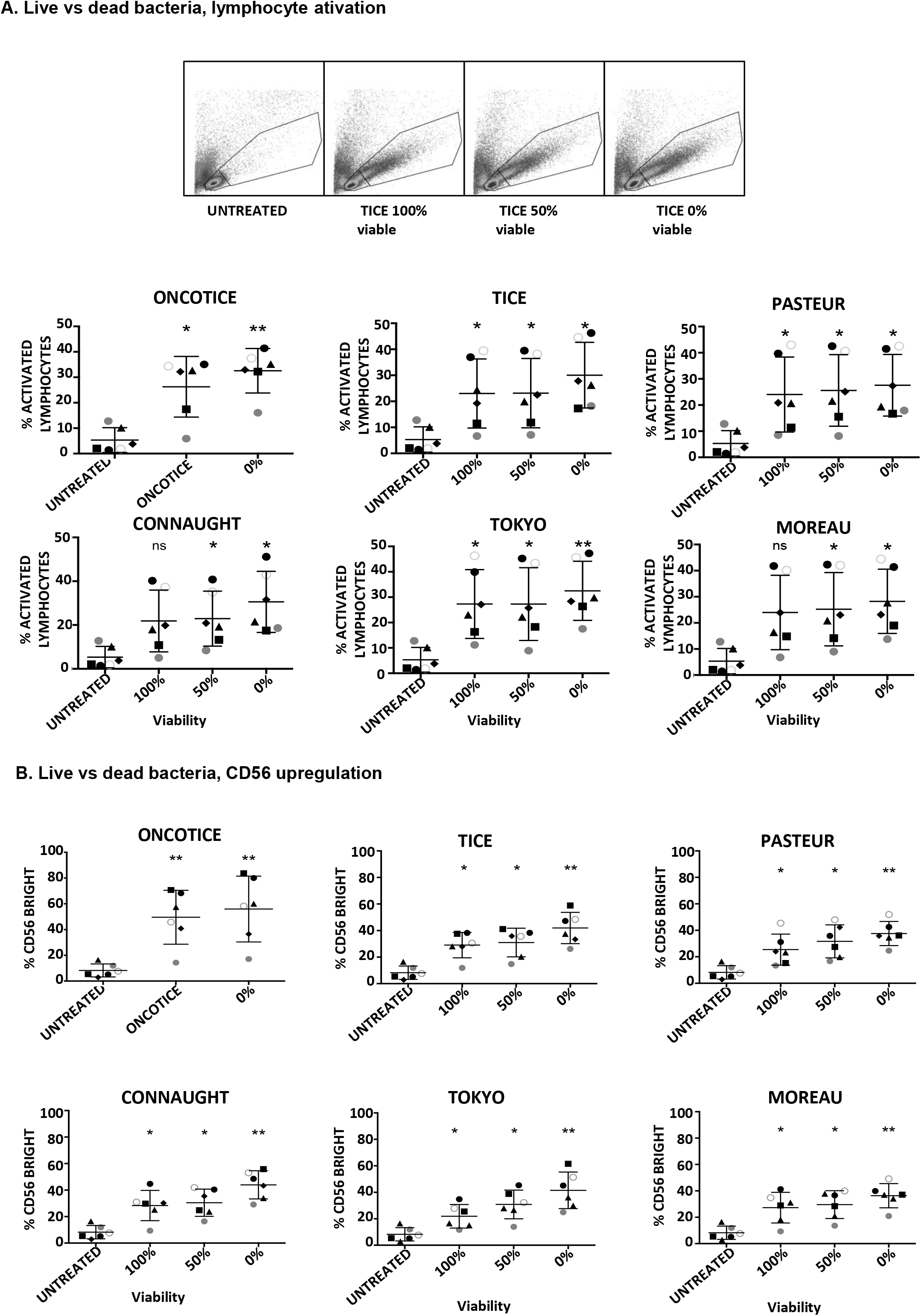
Lymphocyte activation occurs in co-cultures using either live or dead BCG. PBMCs from healthy donors were incubated with the indicated BCG substrains. The ratio was calculated to mimic the amount of total bacteria in the commercial vaccine, 6:1 ratio (total bacteria to PBMC) from cultures either 100% viable, 50% viable or completely inactivated by 20 min at 100°C. At day 7, cells in suspension were recovered from the co-culture, centrifuged and analysed by flow cytometry. **A.** The percentage of activation was determined by identifying the resting and activating lymphocyte regions by FSC vs SSC. The percentage of activated lymphocytes was plotted for each donor (different symbols) and viability condition. Statistical analysis was performed using one-way ANOVA comparing each condition with the untreated culture (*, p<0.05; **, p<0.01; ns, non-significant). **B.** The percentage of the CD56^bright^ population was obtained within the NK cell region for each donor and analysed as in A.

These experiments show that NK cells can be activated when non-viable BCG is co-cultured with PBMCs. To further confirm this hypothesis, the capacity of different laboratory-grown BCG substrains using approximately 10% of live bacteria (proportion found in OncoTICE) to activate PBMCs was tested showing lymphocyte activation once again (Figure 3A, left panel). Although the final amount of different subtypes of lymphocytes varied slightly, all the BCG substrains tested gave rise to the upregulation of CD56 on NK cells (Figure 3A, right panel), resulting in degranulation against bladder cancer cells (Figure 3B). Indeed, upregulation of CD56 was significant for all seven donors tested, compared to untreated PBMCs, and the percentage of CD56^bright^ was very consistent when comparing between the different substrains. In parallel, the capacity of these activated NK cells to degranulate after recognition of two bladder cancer cell lines [T24, a grade 3 cell line recognized by activated NK cells from most donors, and RT112, a low grade cell line usually poorly recognized by activated NK cells, [17]] was tested. These experiments demonstrate an increase in degranulation of CD56^bright^ NK cells when using different BCG substrains containing approximately 10% of viable bacteria (Figure 3B). To sum up, these results suggest that dead mycobacteria in co-culture with PBMCs contribute to the activation of NK cells for an enhanced anti-tumoral activity.

**Figure 3.**
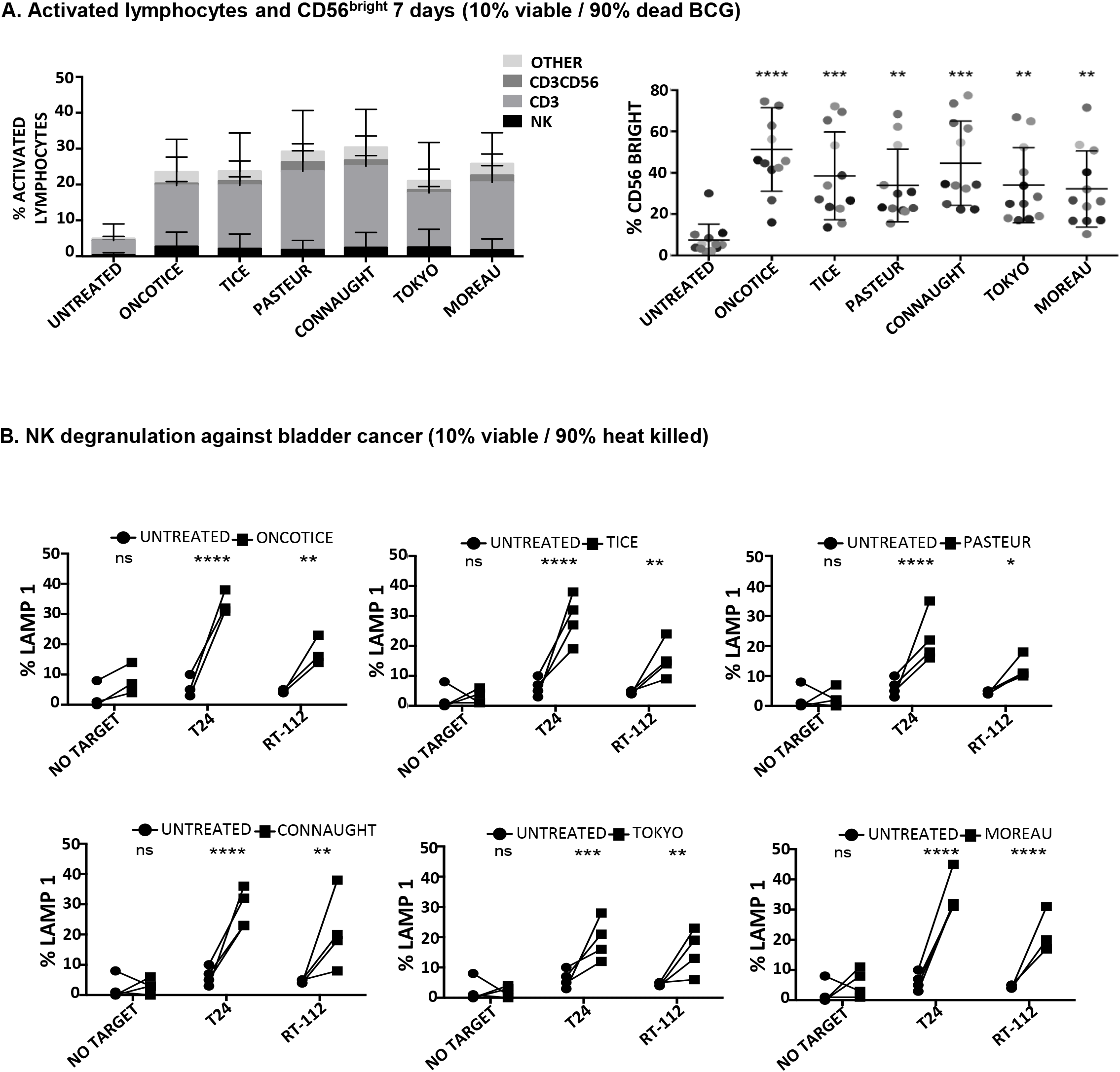
Lymphocyte populations and NK degranulation after co-cultures with 10% viable BCG. PBMCs from healthy donors were incubated with the indicated substrains of BCG at a 1:10 ratio (live bacteria to PBMC), using cultures containing approximately 10% viable bacteria plus 90% inactivated bacteria by 20-30 min at 100°C. OncoTice resuspended from the commercial vial was used as a control. At day 7, cells in suspension were recovered from the co-culture, centrifuged and analysed by flow cytometry. **A.** The percentage of activated lymphocytes was determined by identifying the resting and activating lymphocyte regions by FSC vs SSC and further analysed to distinguish T cells from NK cells by staining with antibodies against population markers (CD3, CD56) (left). The percentage of activated lymphocytes was plotted for each donor (different populations are displayed in different colours). The percentage of the CD56^bright^ population within the NK cell region was obtained for each donor (different colours) and analysed by one-way ANOVA (**, p<0.01; ***, p<0.001; ****, < 0.0001) (right). **B.** Degranulation of NK cells against bladder cancer cells (T24, RT-112) was measured by analysing surface LAMP-1 (CD107a) within the CD3^-^CD56^+^ region. Different donors are represented by symbols and compared with the untreated culture. Statistical analysis was performed using one-way ANOVA (**, p<0.01; ***, p<0.001; ****, p<0.0001; ns, non-significant).

### Different mycobacterial fragments lead to the activation of different immune populations

Since dead BCG contributed to the activation of CD56^bright^ NK cells, experiments were designed to study different PBMC subpopulations. The idea of using fragments of bacteria or different strains (including non-pathogenic) to avoid toxicity in bladder cancer patients has been studied on several occasions [20-22]. Thus, different subcellular fractions of mycobacterial extracts from *M. bovis* and BCG-Tice were used in co-culture experiments with PBMCs. *M. bovis* (Strain AF 2122/97ATCC^®^ BAA-935^™^) and its fractions were obtained from BEI Resources, NIAID, NIH; BCG-Tice fractions were prepared from BCG-TICE cultures by extraction with organic solvents (see Methods). Whole irradiated *M. bovis*, and the corresponding fractions, membrane, cytosolic, total lipids and individual lipids [(Trehalose-6,6-dimycolate (TDM) and mycolic acid] were incubated at different concentrations in co-culture with PBMCs (7 days) and the percentage of activated lymphocytes and NK subpopulations was analyzed by flow cytometry (Supplementary Figure 3A). Since most of the compounds tested could activate lymphocytes, experiments to define the relative activation of each lymphocyte subpopulation were performed (Figure 4A). Whole irradiated *M. bovis* and membrane fractions led to an increase in lymphocyte activation and, although an increase in CD56^bright^ cells could be observed, the population of NK cells did not proliferate as much as that of CD3 lymphocytes. In particular, CD4 cells were significantly expanded in cultures where the membrane fraction was added (Figure 4B). In agreement with these findings, low degranulation capacity of NK cells against the two bladder cancer cell lines, T24 and RT112, was found in these cultures (Figure 4C). These experiments demonstrated that, although OncoTICE led to a lower percentage of activated lymphocytes than *M. bovis*, the composition of activated subpopulations was more skewed towards NK cells in the former leading to higher percentage of CD56^bright^ upregulation and NK degranulation capacity. On the other hand, CD4 cells were activated more efficiently by *M. bovis* fragments or irradiated cells. The cytosolic fraction of *M. bovis* only could stimulate NK degranulation against T24 cell line and a non-significant upward trend could be observed for RT112. The lipidic fractions of *M. bovis*, in the concentrations used for these experiments, did not stimulate the PBMC culture.

**Figure 4.**
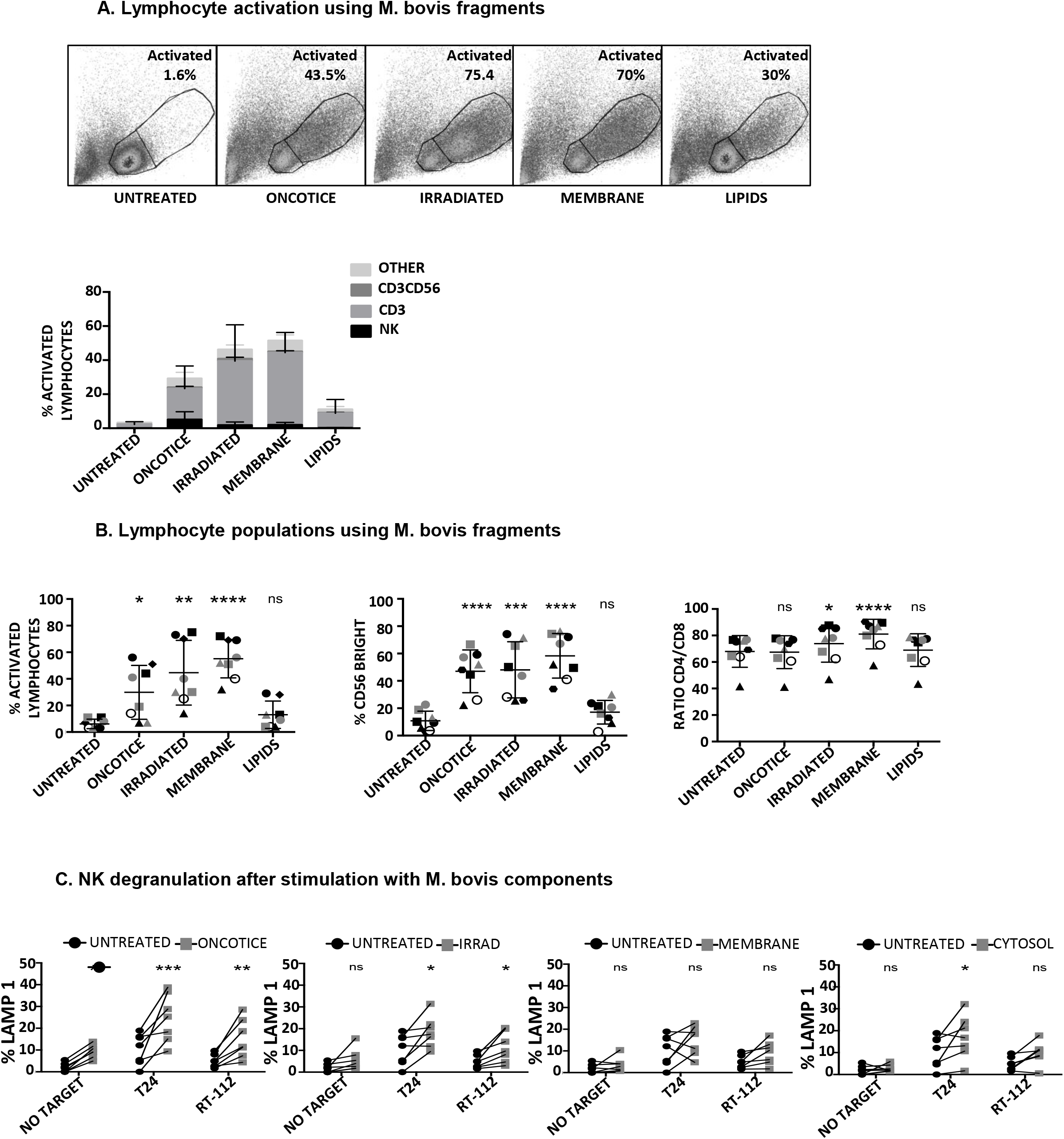
Lymphocyte activation using *M. bovis* and its subcellular fractions. PBMCs from healthy donors were incubated with the indicated fractions of *M. bovis* obtained through BEI Resources, NIAID, NIH. At day 7, cells in suspension were recovered from the co-culture, centrifuged and analysed by flow cytometry. **A.** The percentage of activation was determined by identifying the resting and activating lymphocyte regions by FSC vs SSC and further analysed to distinguish T cells from NK cells by staining with antibodies against population markers (CD3, CD56) (top). The percentage of activated lymphocytes was plotted for each donor (different populations are displayed in different shades of grey/black) (bottom). **B.** The percentage of the CD56^bright^ population was obtained within the NK cell region and the CD4/CD8 ratio was obtained within the CD3^+^ region. Data are displayed as different symbols for each donor. Statistical analysis was performed using one-way ANOVA comparing each condition with the untreated culture (*, p<0.05; **, p<0.01; ***, p<0.001; ****, p<0.0001; ns, non-significant). **C.** Degranulation of NK cells against bladder cancer cells (T24, RT-112,) was measured by analysing surface LAMP-1 (CD107a) within the CD3^-^CD56^+^ region. Different donors are represented by symbols and compared with the untreated culture. Statistical analysis was performed using one-way ANOVA (*, p<0.05; **, p<0.01; ***, p<0.001; ns, non-significant).

The same type of experiments was performed to evaluate NK activation using BCG-TICE fragments. (Supplementary Figure 3B; Figure 5). In this case, delipidated-BCG, as well as the lipidic and the aqueous fractions were used. In contrast to *M. bovis* fragments, TICE lipids, mildly activated PBMC cultures while the aqueous phase did not activate lymphocytes at all (Figure 5A). The more activated cultures were obtained upon incubation with delipidated BCG. No fragment was as efficient as OncoTICE at stimulating NK CD56 upregulation, however the percentage of activated lymphocytes was higher in cultures with both delipidated-BCG and lipid fragments than in untreated PBMCs (Figure 5B). Both the lipid and the aqueous fraction led to a low level of CD56^bright^ activation, but when present, these cells were able to recognize bladder cancer cells in degranulation assays (Figure 5C). In fact, the amount of granzyme B and perforin in CD56^bright^ NK cells was clearly increased when PBMCs were treated with delipidated-BCG in a comparable manner to cultures stimulated with BCG-Tice (Figure 6). These data suggest that different mycobacteria fractions can activate immune populations differently and that, depending on the intended use of BCG, different fragments could be enough to produce the desired effect.

**Figure 5.**
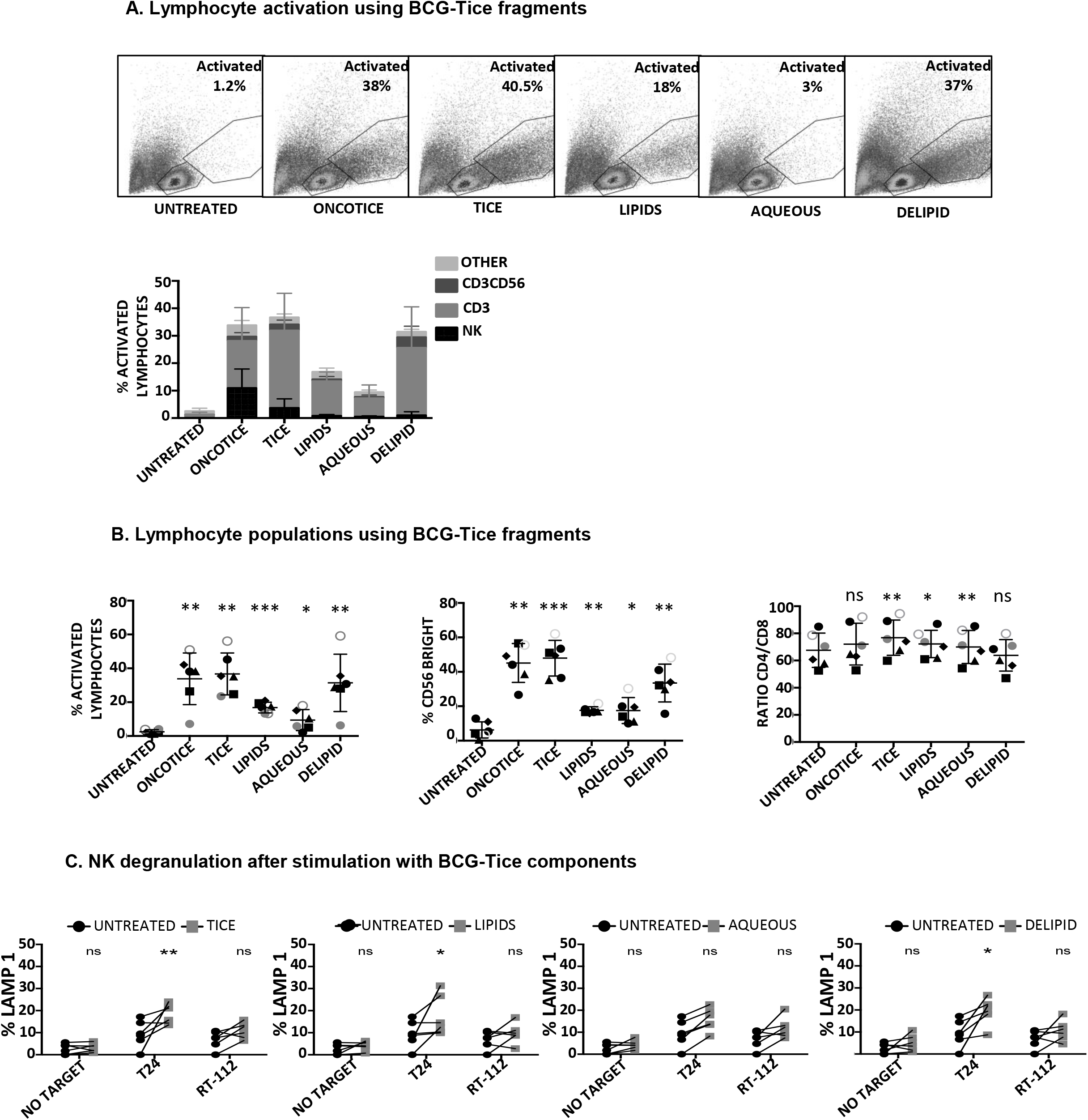
Lymphocyte activation using Tice BCG subcellular fractions. PBMCs from healthy donors were incubated with the indicated fractions of Tice BCG (see methods). At day 7, cells in suspension were recovered from the co-culture, centrifuged and analysed by flow cytometry. **A.** The percentage of activation was determined by identifying the resting and activating lymphocyte regions by FSC vs SSC and further analysed to distinguish T cells from NK cells by staining with antibodies against population markers (CD3, CD56). The percentage of activated lymphocytes was plotted for each donor (different populations are displayed in different shades of grey/black) (bottom). **B.** The percentage of the CD56^bright^ population was obtained within the NK cell region and the CD4/CD8 ratio was obtained within the CD3^+^ region. Data are displayed as different symbols for each donor. Statistical analysis was performed using one-way ANOVA comparing each condition with the untreated culture (*, p<0.05; **, p<0.01; ***, p<0.001; ns, non-significant). **C.** Degranulation of NK cells against bladder cancer cells (T24, RT-112,) was measured by analysing surface LAMP-1 (CD107a) within the CD3^-^CD56^+^ region. Different donors are represented by symbols and compared with the untreated culture. Statistical analysis was performed using one-way ANOVA (*, p<0.05; **, p<0.01; ns, non-significant).

**Figure 6.**
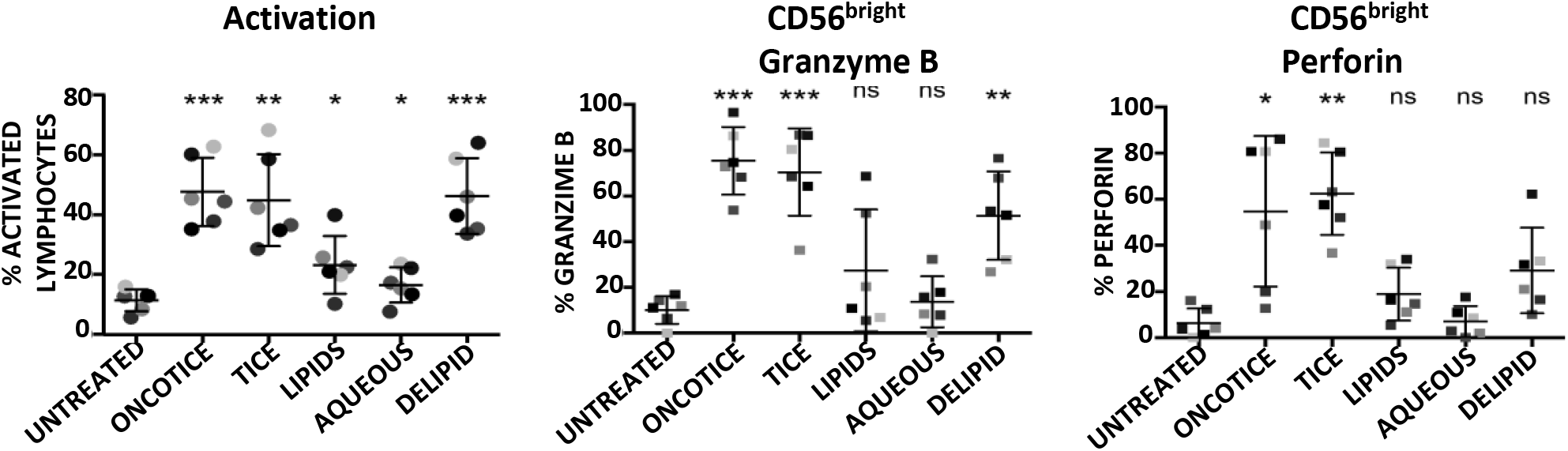
Granzyme B and perforin within NK cells activated using Tice BCG subcellular fractions. PBMCs from healthy donors were incubated with the indicated fractions of BCG-Tice (see methods). At day 7, cells in suspension were recovered from the co-culture, centrifuged and analysed by flow cytometry. The percentage of activation was determined by identifying the resting and activating lymphocyte regions by FSC vs SSC and further analysed to distinguish T cells from NK cells by staining with antibodies against population markers (CD3, CD56). The percentage of activated lymphocytes was plotted for each donor (different colours) (left). The percentage of the CD56^bright^ population was obtained within the NK cell region and the content of Granzyme B (middle) and perforin (right) were determined by intracellular staining. Statistical analysis was performed using one-way ANOVA (*, p<0.05; **, p<0.01; ***, p<0.001; ns, non-significant) and compared with the untreated culture.

## DISCUSSION

BCG has been used both as TB vaccine and to treat bladder cancer for many decades, however, the heterogeneity of action due to the existence of many different substrains still poses many unresolved questions. This study evaluates the effect of BCG heterogeneity in the immune response exerted by NK cells against bladder cancer cells, as a model of the successful immunotherapy used routinely in NMIBC patients. The data presented here confirm that the different mycobacterial strains have different capacities to stimulate leukocytes and that the subpopulations of lymphocytes stimulated by different bacterial compounds diverge.

A key discovery reported here is that, although all BCG substrains can stimulate NK cell proliferation and response against high grade bladder cancer, clearly some substrains provoke NK cell degranulation more strongly than other. Different BCG substrains were generated from 1921 when the original culture was continuously grown separately in different countries [7,11]. Several genetic groups of BCG appeared subsequently differing in regions of difference (RD) that have particular deletions, insertions and mutations. RD1 deletion was the first main genetic change that occurred between *M. bovis* and BCG. Interestingly, in our degranulation experiments, the stronger increase in response against T24 bladder cells occurred using as stimulation of PBMCs Tice, Connaught and Moreau. The genetic branch of the Moreau substrain separated early, in 1924, before the loss of IS6110, and belongs to the same group as Tokyo, 1925, which resulted in a more discrete NK response. Moreau and Tokyo differ in the regions, RD16 and ΔRv3405c, while Tice (1934) and Connaught (1948) belong to the same genetic group [tandem duplication 2 (DU2) group 4] and their main difference is the loss of RD15 that occurred in Connaught. Whether these genes are involved in the production of a stronger or milder NK cytotoxic response was not explored here, but could be the basis of the functional differences found among the different BCG substrains.

A second finding from this work is that vaccine preparations contain, in addition to the CFU specified, a large number of dead bacteria and this prompted a series of experiments to evaluate the effect of dead mycobacteria on stimulation of leukocytes. Here, we confirm that the capacity of dead bacteria to stimulate lymphocyte proliferation and NK cell activation in co-cultures of PBMCs was similar to the stimulation caused by live bacteria. In certain strains, dead bacteria were even more potent. Early animal model experiments already concluded that dead BCG could mediate an anti-tumour response when bacteria were applied by intradermal injection [23] and, since then, several approaches have been investigated in order to eliminate live bacteria in bladder cancer treatment to avoid toxicity, for example using bacterial cell wall and nucleic acids that could benefit patients failing other therapies [24]. This study provides a first evaluation of the impact of different mycobacterial components in the activation of NK cells to recognise bladder cancer cells. The experiments using *M. bovis* and BCG-Tice subcellular components confirmed that CD56^bright^ NK cells respond strongly against bladder cancer and that this activation might be modulated by different mycobacterial compounds, but again, experiments showed differences between the two mycobacteria strains. While the lipidic fraction of *M. bovis* did not stimulate any immune population, the lipids from Tice activated mainly CD3, increasing the ratio CD4/CD8. On the other hand, delipidated BCG-Tice did not change the proportion of CD4/CD8, but caused the increase of effector cells both CD3^+^CD56^+^ and NK cells. All these data support the idea that new inactivated BCG-based preparations could be possible both for cancer treatment and other situations in which NK cell stimulation could be beneficial. In fact, new approaches using intravenous BCG to stimulate certain immune populations, such as Th1/Th17 CD4 or CD8 T cells, proposed to increase TB protection in youngster or adults could benefit from the use of fractions and not viable bacteria [25]. Our study proposes that a particular immune population could be stimulated choosing the right BCG substrain or component, and thus, more research should be conducted to comprehensively identify the immune cells subpopulations responding to specific mycobacterial products. The use of NK cells as a therapeutic agent in cancer has been long discussed and nowadays, a large body of research is available on the use of NK cells for treatment of cancer, both autologous or allogeneic after ex vivo stimulation. Recent research and clinical trials are starting to generate promising results [26-29]. The work presented here could open a new manner for NK stimulation with therapeutic purposes.

In conclusion, here we demonstrate that BCG stimulates the activation of NK cells through the interaction with mycobacteria subcellular components and this type of stimulation could be of interest beyond bladder cancer treatment. The presence of dead mycobacteria in lyophilised BCG preparations suggest that live bacilli could be less important than initially thought for cancer elimination and this opens new possibilities for a desirable decrease of unwanted effects caused by live BCG.

## Acknowledgements

This work was supported by grants from the Spanish Ministry of Science and Innovation (RTC-2017-6379-1, RTI2018-093569-B-I00 (MCIU/AEI/FEDER, EU) (MVG) and the regional government of Madrid (S2017/BMD-3733-2); Generalitat of Catalunya (2017SGR-229) (EJ). The group of MVG belongs to the research network TENTACLES (RED2018-102411-T) funded by the Spanish Ministry of Science.

The authors would like to thank Dr Sylvia Leao, for the gift of BCG. The following reagents were obtained through BEI Resources, NIAID, NIH: γ-irradiated whole cells *Mycobacterium bovis*, Strain AF 2122/97 (ATCC^®^ BAA-935^™^) Catalog No. NR-31210; Total Lipids, Catalog No. NR-44100, Cell Membrane Fraction, Catalog No. NR-31214 and Cytosol Fraction, Catalog No. NR-31215 from *M. bovis;* Trehalose-6,6-dimycolate (TDM), Catalog No. NR-14844and Mycolic Acid, Catalog No. NR-14854 from *Mycobacterium tuberculosis*, Strain H37Rv.

## Author contributions

GE, NA, acquired, analysed and interpreted data

EJ, OA, MMH, generated reagents

NA, CM, MVG, provided material support and supervised the study

All authors, participated in drafting and critical revisions of the article

---

**Supplementary Figure 1.**
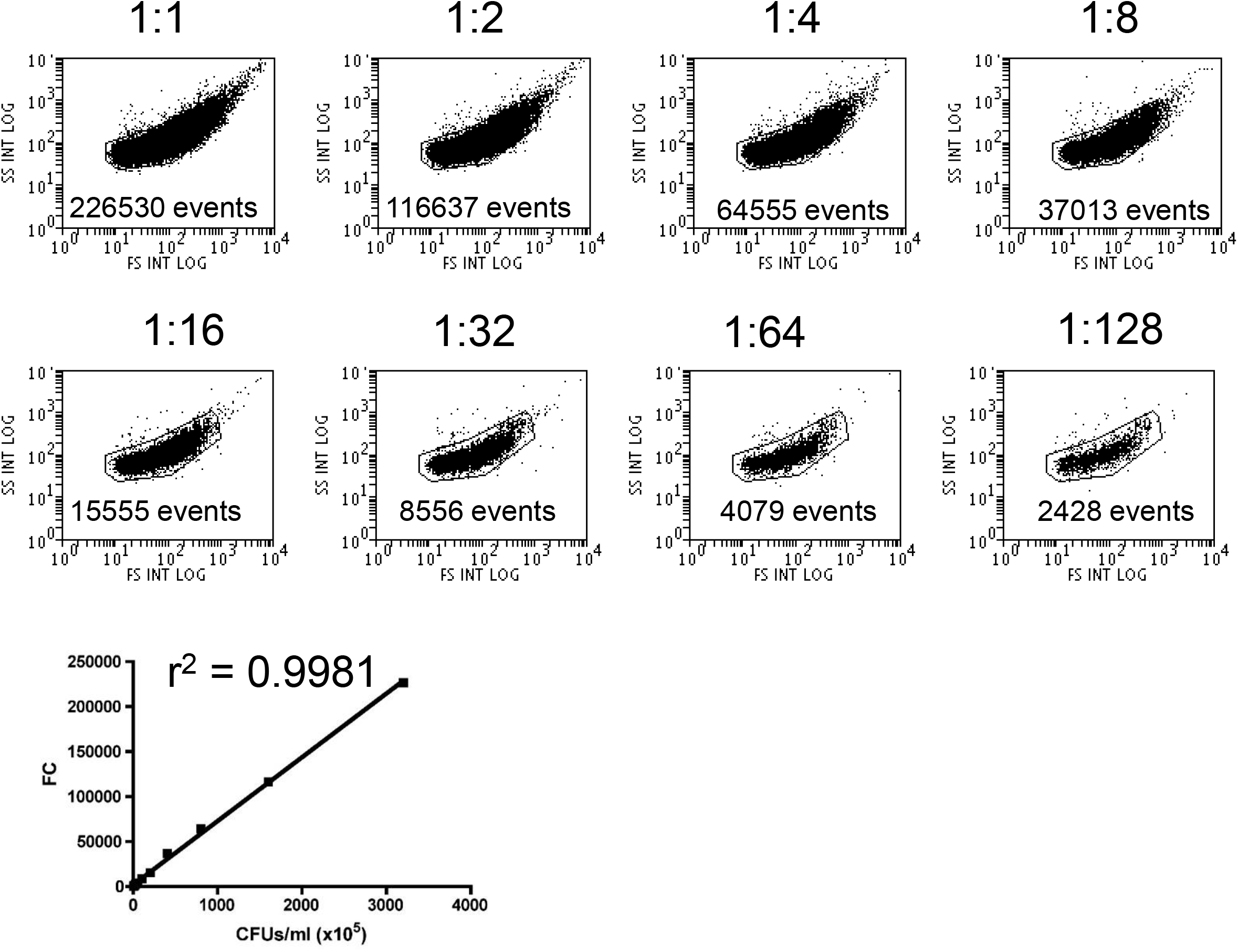
The number of BCG cells estimated by flow cytometry. Serial dilutions, as indicated, of BCG grown in Middlebrook 7H10 medium were prepared and analysed by flow cytometry at constant speed during 30 seconds. The number of events within the FSC vs SSC region corresponding to bacteria (A) were used to build a calibration curve (B).

**Supplementary Figure 2.**
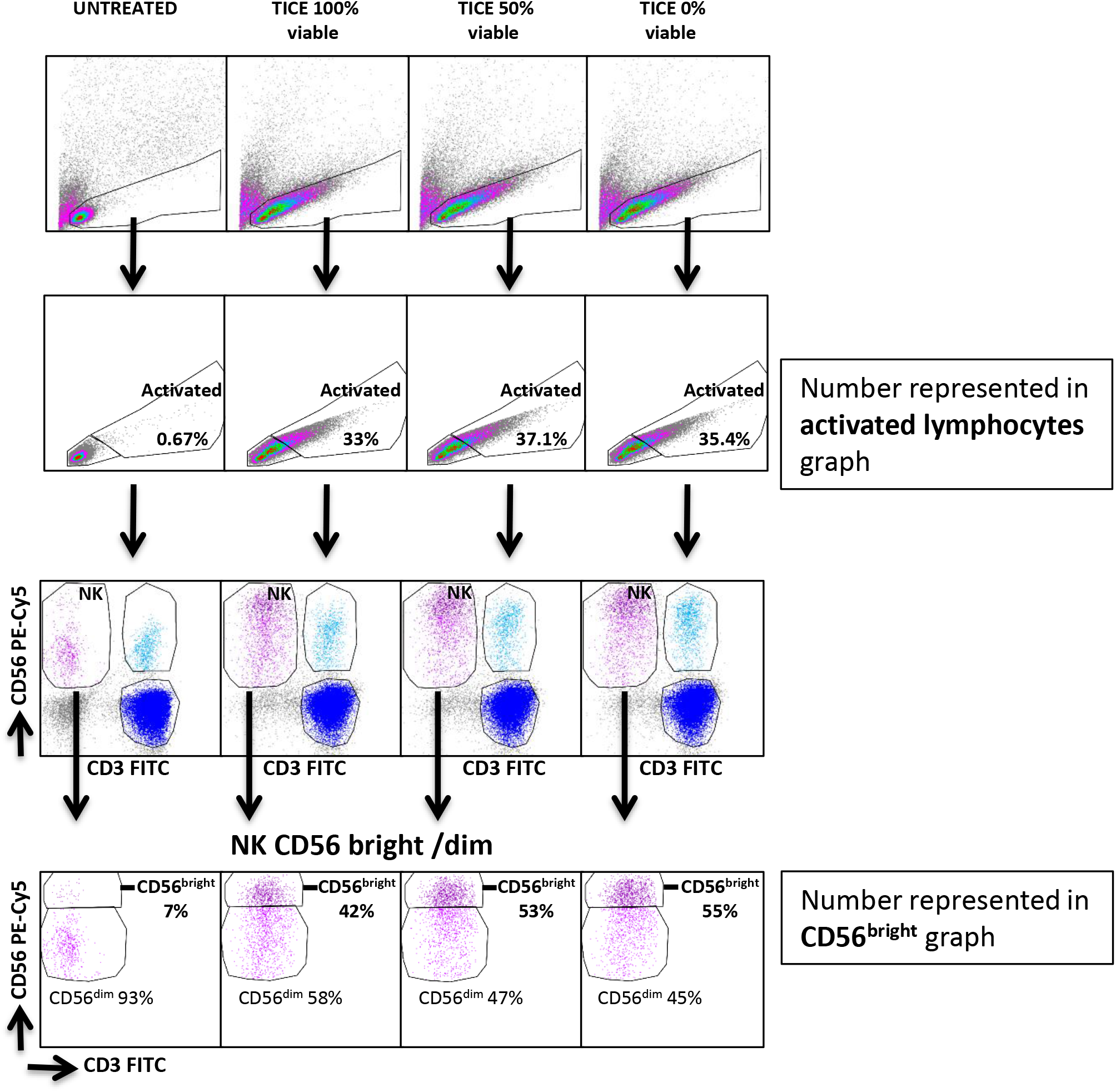
Gating strategy for activated lymphocytes and NK CD56^bright^ population after co-culture with BCG. PBMCs from a healthy donor were incubated with the indicated concentrations of BCG-Tice at a 6:1 ratio (bacteria to PBMC). At day 7, cells in suspension were recovered from the coculture, centrifuged and analysed by flow cytometry. Once dead cells were eliminated, the percentage of activated lymphocytes was determined by identifying the resting and activating lymphocyte regions in the FSC vs SSC plot. The percentage of activated lymphocytes from this plot was included in the subsequent graphs. NK cells were selected within the whole lymphocyte population in a CD3 vs CD56 plot. The percentage of CD56^bright^ population, within the NK cell region, was plotted.

**Supplementary Figure 3.**
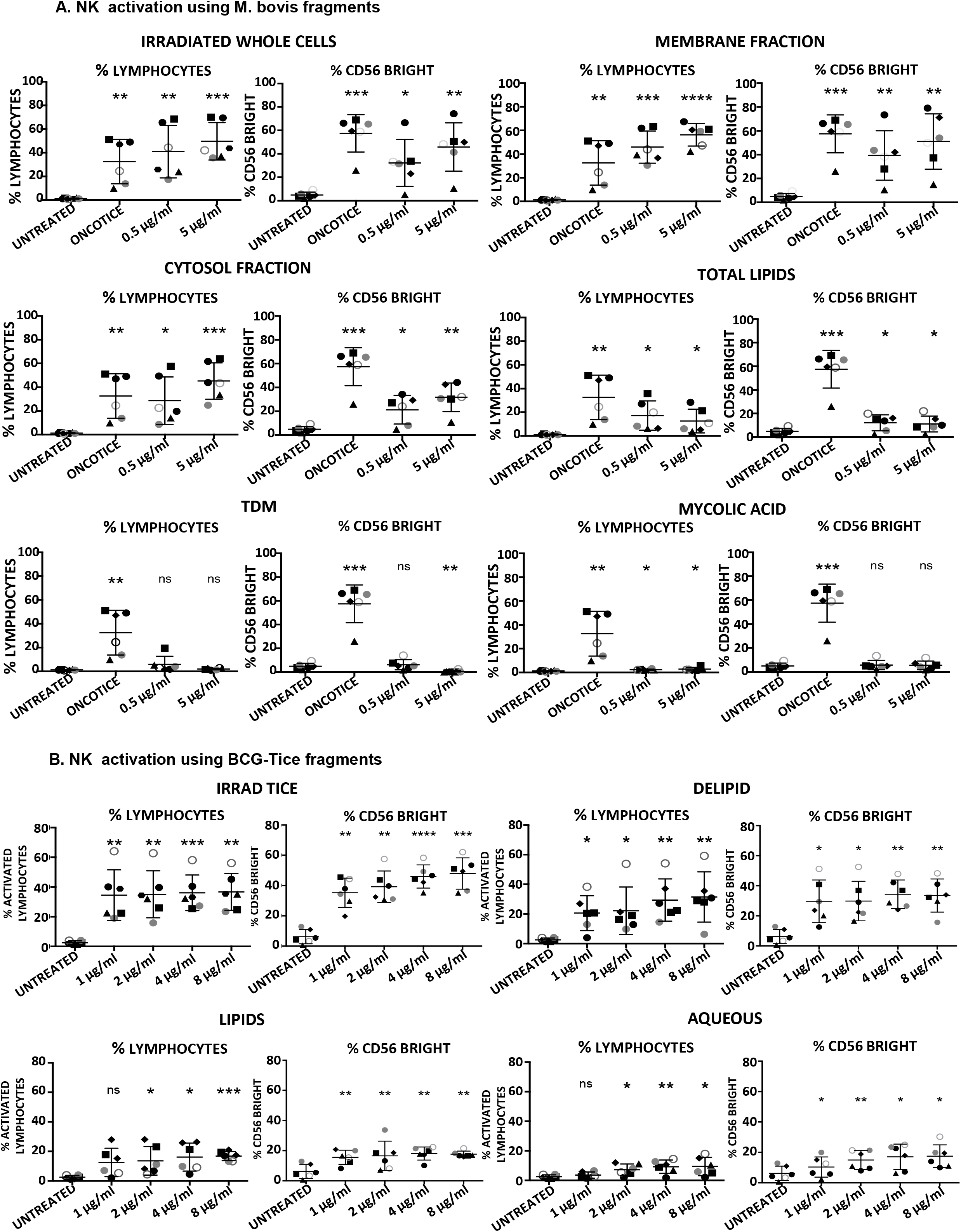
Lymphocyte activation after co-culture with mycobacterial subcellular fragments. **(A) *M. bovis.* (B) BCG-TICE**. PBMCs from healthy donors were incubated with the indicated concentrations of mycobacterial fractions (see methods). OncoTice was used as control at a 1:10 ratio (bacteria to PBMC). At day 7, cells in suspension were recovered from the co-culture, centrifuged and analysed by flow cytometry. The percentage of activated lymphocytes was determined by identifying the resting and activating lymphocyte regions by FSC vs SSC and further analysed to distinguish NK cells by staining with antibodies against population markers (CD3, CD56). The percentage of activated lymphocytes (left) and the percentage of the CD56^bright^ population within the NK cell region (right) were plotted for each donor (different symbols). Statistical analysis was performed by one-way ANOVA (*, p<0.05; **, p<0.01; ***, p<0.001; ****, < 0.0001; ns, non-significant).

